# Histo-morphological development of the digestive tract of larval hybrid Malaysian mahseer (*Barbonymus gonionotus* ♀ × *Tor tambroides ♂*)

**DOI:** 10.1101/2020.06.19.161976

**Authors:** Mohd Salleh Kamarudin, Muhammad Azfar Ismail, Fadhil Syukri, Kamil Latif

**Affiliations:** Department of Aquaculture, Faculty of Agriculture, Universiti Putra Malaysia, 43400 Serdang, Selangor, Malaysia; International Institute of Aquaculture and Aquatic Sciences, Universiti Putra Malaysia, Batu 7, Jalan Kemang 6, Teluk Kemang, Si Rusa, 71050 Port Dickson, Negeri Sembilan, Malaysia; Department of Animal Science and Fishery, Universiti Putra Malaysia Bintulu Campus, 97008 Bintulu Sarawak, Malaysia

**Keywords:** carp, digestive capability, histology, weaning, gut index

## Abstract

New carp hybrids are being developed for the aquaculture industry to support rising seafood demands. The present study was carried out to observe the changes in digestive tract, histology and functional capabilities of the new hybrid carp larvae for a better understanding of its digestive capability and the prediction of its best weaning time to a compound diet. A tubal digestive tract was elongated to the anus and buccal cavity by 3 DAH that coincided with the mouth opening and the start of exogenous feeding. A functional stomach was observed at 7 DAH with the relative gut index (RGI) of 10.7 ± 0.06. A layer of supranuclear protein was observed with lipoprotein at the outer layer of the digestive tract at 7 DAH. The morpho-histological results of this study indicated that hybrid Malaysian mahseer larvae should be able to digest, ingest and absorb an artificial diet beginning from 7 DAH. At this stage, the hybrid larvae could be gradually or perhaps totally weaned to an artificial diet of a suitable particle size.

## 1. Introduction

In general, the morphological and histological studies of larval fish digestive tract have been conducted to identify the suitable time or period to wean the larvae from live foods to an artificial diet and types of feeding required by fish larvae (Avila and Juario, 1987; Cousin et al., 1987; Ferraris et al., 1987; Munilla-Moran and Stark, 1989; Walford et al., 1991). Digestive tract comprises the special morphological characteristics and adaptations for food and feeding protocols (Kapoor et al., 1976). A better understanding of digestive tract leads to the provision of favorable foods and right weaning period to an artificial diet during the larval rearing. There are several activities occur in a digestive system including absorption, secretion, and cellular adhesion to converting the foods into nutritional components for growth, repair, and others (Smith, 1980).

The digestive tract length and features can also indicate the types of fish feeding behavior such as detritivores, herbivores, omnivores, carnivores and frugivores (Kramer and Bryant, 1995). Optimum feeding and rearing protocol can also be developed from the information on the functional capabilities of the digestive tract and nutritional physiology of larval fish (Segner et al., 1993; Walford and Lam, 1993). Many studies have been conducted in teleosts to observe the digestive tract development and weaning time during larval and juvenile stages of fish (Baglole et al., 1997; Gordon and Hecht, 2002; Loewe and Eckmann, 1988; Mai et al., 2005; Ribeiro et al., 1999). The digestive tract of fish larvae is less coiled in comparison to adults, and the digestive glands and enzyme production are usually not fully developed in some fish during the weaning period (Holt, 1992).

In general, an artificial diet is gradually introduced once the fish larvae have reached a suitable weaning time which is usually coincided the full development of the stomach. Live food organisms such as rotifers, microworms, artemia and moina are the common food given to the larvae as moving preys readily attract larvae. However, the production of most live feed is expensive and unpredictable with variations in quality (Cahu and Zambonino Infante, 2001; Dabrowski, 1989) that necessitates a shift to an artificial and complete diet as earlier as possible (Jones et al., 1993). In some cases, the use of artificial diets produce superior results in comparison to live feeds. Larval African catfish, *Clarias gariepinus* has a similar growth performance when fed with an artificial diet but with a lower survival rate compared to those fed with live food (Kamarudin et al., 1996). The objective of this study was to observe the morphological and histological development of digestive tract of hybrid Malaysian mahseer larvae. The study could provide useful information for determining the best feeding protocol of hybrid Malaysian mahseer larvae.

*Tor tambroides* is one of the most expensive freshwater fish in South East Asia and favoured species in aquaculture as well as sport fishing (Ng, 2004). However, the commercial breeding of *T. tambroides* faces several challenges due to its partial spawning, low survival and fry quality (Ingram et al., 2005). The earlier success of crossbreeding lemon fin barb and silver barb, and the aquaculture of its hybrid (DOF, 2012) had motivated researchers in the Universiti Putra Malaysia to crossbreed *T. tambroides* male and *Barbonymus gonionotus* female with the intention to reduce the pressure on the fisheries of *T. tambroides* and the collection of wild fries for its aquaculture. *B. gonionotus* is a well-established freshwater aquaculture species that is known for its good taste, fast growth rate and all year around spawning (Basak et al., 2014; Sarker et al., 2002). The first successful induced breeding of hybrid Malaysian mahseer was achieved in 2017. Research is being conducted the researchers to ascertain its optimum culture conditions and increase the hybrid hatching rate from the current very low 5%. Despite having a low hatching rate, the survival of larvae is high at about 90% and higher (Azfar Ismail et al., 2018).

## 2. Materials and methods

This study was carried out at the Department of Aquaculture, Faculty of Agriculture UPM. Broodstock of silver barb females (0.7-1.5 kg) and Malaysian mahseer males (2-3 kgs) were obtained from a fish farmer in Selangor, Malaysia. The fish were injected intramuscularly with a commercial Ovatide® hormone at a dosage of 0.4 ml kg^-1^ for females and 0.2 ml kg^-1^ for males. The eggs were stripped from the silver barb females 5-6 h after the hormone injection and mixed with the sperm of the Malaysian mahseers. Fertilised eggs were distributed into three 75 L incubation glass aquaria at 1620 eggs l^-1^ for hatching. The eggs started to hatch as early as 13 h after fertilization.

Newly hatched healthy larvae were carefully collected using 3 ml of disposable pipette and distributed into another three 75 L glass aquaria at 10 larvae l^-1^ with a gentle aeration provided. Microworms (*Panagrellus redivivus*) with the size of 50-60 µm were introduced for the first feeding of the larvae at 3 days after hatching (DAH). By 5 DAH, the larvae were fed with newly hatched artemia nauplii with the size of 180-250 µm and then gradually weaned to a combination of nauplii and a formulated microdiet (Larval AP100, Ziegler Bros USA) at 8 DAH. Water quality was maintained by changing 30% of rearing media twice a week.

Twenty fish larvae were sampled daily randomly from 1 to 10 DAH and thereafter at 2-day intervals until 23 DAH. Ten fish larvae were observed under a light microscope (Zeiss Primo Star) fitted with a microscope eyepiece camera (Dino-eye). Morphometric data (total length (TL), body weight (BW), gut length (GL)] were recorded. The drawings of digestive tract development were made from the images generated through the microscope. The relative gut index (RGI) of each larva was estimated using the following formula:

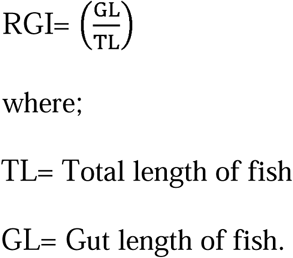

Ten larvae were also sampled for general histological examination of the digestive tract structure. Samples were fixed in Bouin’s solution for 8-12 hours and preserved in 70% ethanol. Samples underwent tissue processing through dehydration, clearing and infiltration. Dehydration was done at 70, 80, 95 and 100% ethanol and followed by a clearing using xylene and infiltration using molten wax (Thermo Scientific Microm STP 120). Samples were subsequently put into cassettes and molten wax using Thermo Scientific Histostar. Samples were cut into 6 µm sections (Thermo Scientific Microm HM 340E) and placed on microscopic slides on a slide drying bench. The slides were then stained with haematoxylin-eosin for the coloration of the tissues and muscles. Once completed, the slides were observed under light microscope (Zeiss Primo Star) fitted with a microscope eyepiece camera (Dino-eye).

## 3. Results

Table 1 shows the morphometric measurements [TL, GL, and RGI] of hybrid Malaysian mahseer larvae. A strong polynomial relationship GL and DAH (GL = 0.0796DAH^2^ + 0.7465DAH + 0.0554 r = 0.994) was observed (Figure 1). RGI values were less than 1.0 at early stage and increased as the fish grew starting from 7 DAH (10.7 ± 0.06 of RGI). The digestive tract length at 23 DAH was longer from the fish total length with the RGI value was 2.54 ± 0.11.

**Table 1.** Morphometric measurement of Malaysian mahseer hybrid in larval stage in total length (TL), gut length (GL), and relative gut index (RGI).

| Larval age | TL (mm) | GL (mm) | RGI |
| --- | --- | --- | --- |
| 1 DAH | 4.06 ± 0.27 | 1.84± 0.15 | 0.46 ± 0.06 |
| 2 DAH | 4.59 ± 0.15 | 2.65± 0.14 | 0.58 ± 0.03 |
| 3 DAH | 4.80 ± 0.14 | 3.01± 0.15 | 0.63 ± 0.04 |
| 4 DAH | 5.01 ± 0.16 | 3.98± 0.36 | 0.80 ± 0.08 |
| 5 DAH | 5.85 ± 0.17 | 5.05± 0.28 | 0.86 ± 0.05 |
| 6 DAH | 7.17 ± 0.15 | 6.52± 0.38 | 0.91 ± 0.05 |
| 7 DAH | 7.61 ± 0.20 | 8.17± 0.33 | 1.07 ± 0.06 |
| 9 DAH | $8.34 \pm 0.24$ | $13.17 \pm 0.34$ | $1.58 \pm 0.05$ |
| 11 DAH | $9.95 \pm 0.17$ | $17.48 \pm 0.47$ | $1.76 \pm 0.05$ |
| 13 DAH | $10.98 \pm 0.26$ | $22.61 \pm 0.79$ | $2.06 \pm 0.07$ |
| 15 DAH | $14.17 \pm 0.37$ | $30.68 \pm 0.52$ | $2.17 \pm 0.05$ |
| 17 DAH | $15.63 \pm 0.39$ | $39.05 \pm 0.77$ | $2.50 \pm 0.06$ |
| 19 DAH | $16.81 \pm 0.35$ | $42.71 \pm 1.22$ | $2.54 \pm 0.08$ |
| 21 DAH | $18.90 \pm 0.48$ | $48.19 \pm 1.27$ | $2.55 \pm 0.11$ |
| 23 DAH | $23.56 \pm 0.91$ | $59.81 \pm 0.95$ | $2.54 \pm 0.11$ |

**Figure 1.**
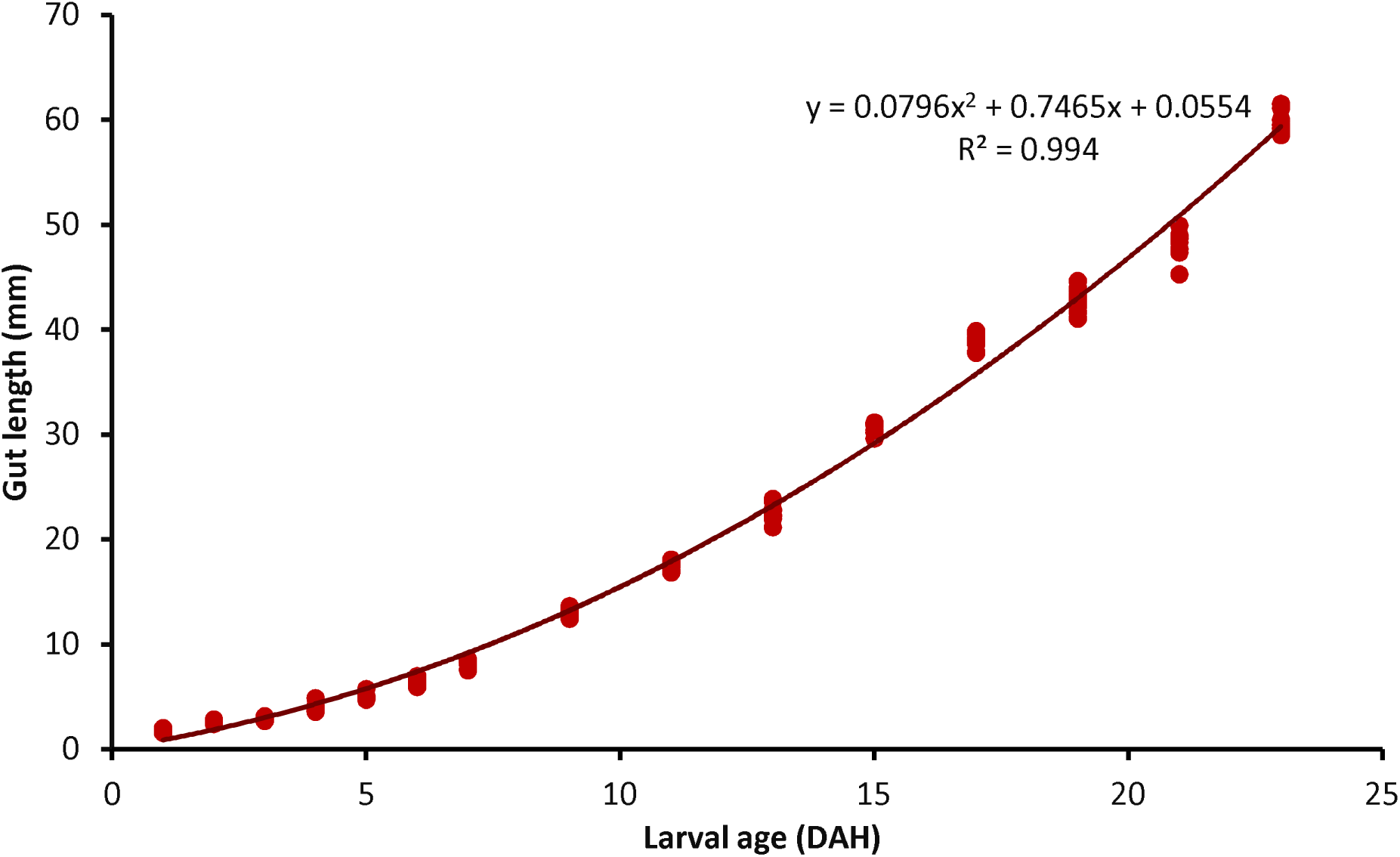
Polynomial relationship between gut length (mm) and larval age (DAH) of Malaysian mahseer hybrid larvae.

The changes in the digestive tract of fish larvae were summarised in Figure 2. At 1-2 DAH, the digestive tract was made of a simple, short, slender and straight tube (1.84 ± 0.15 to 2.65 ± 0.14 mm long) with an undefined stomach bulge and dorsally attached to the large bi-lobed yolk sac (Figure 3a). It was not connected to both mouth and anal openings which were still absent at this stage. The pharynx begins to differentiate from the buccal cavity from 2 DAH together with the presence of simple muscle cells along the region. The digestive tract began to elongate at the oesophagus with the formation of simple muscle cells at 2 DAH (Figure 3b). The oesophagus which was composed of connective tissues and a thin serosa layer with the attachment of goblet cells continued to differentiate with simple cuboidal cells. The goblet cells formed during early larval age were most obvious in the digestive tract from 2 DAH and were clearly visible by 3 DAH (Figure 3c).

**Figure 2.**
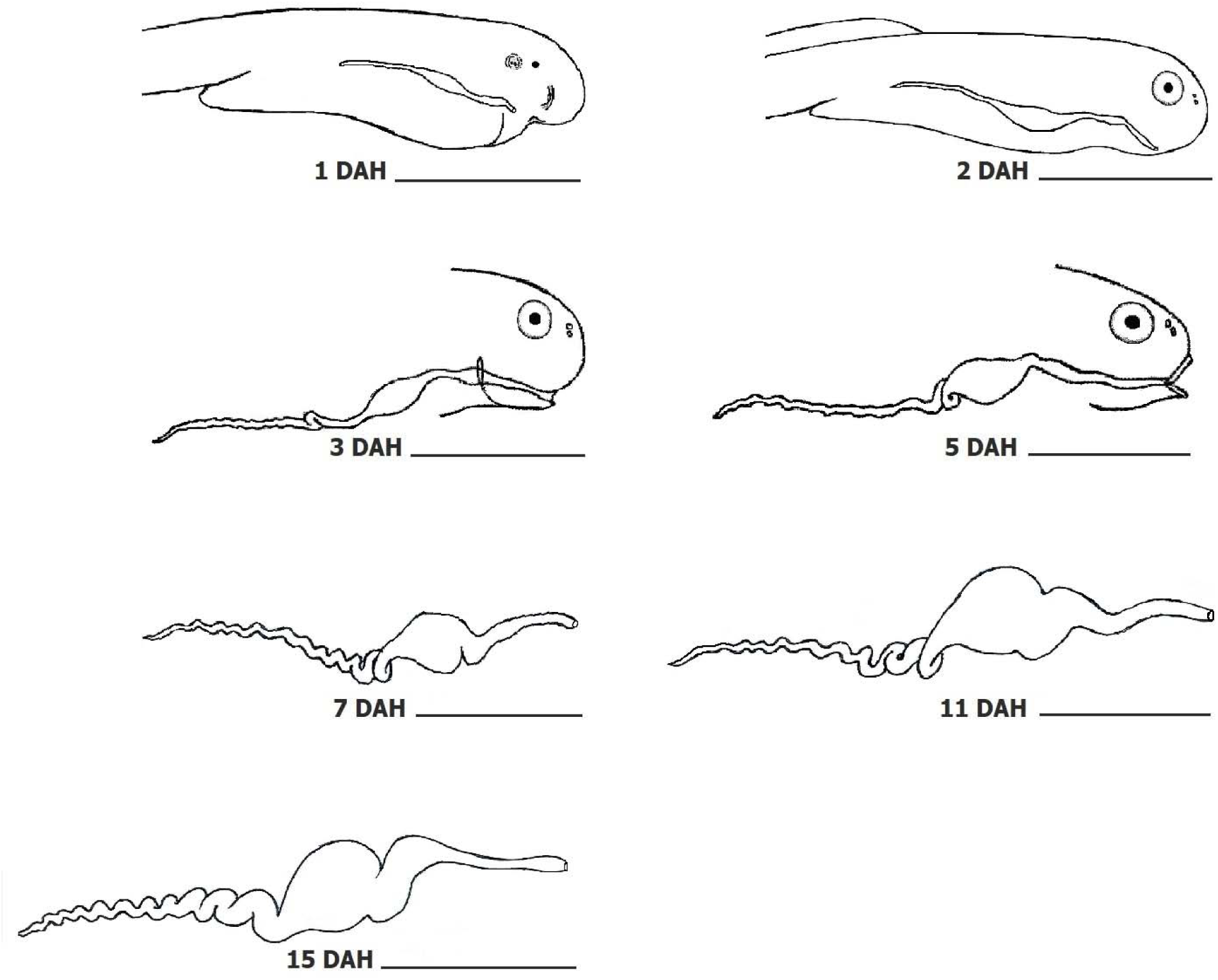
Morphological development of the digestive tract of hybrid Malaysian mahseer from 1 to 15 DAH. No morphological changes in the digestive tract was observed after 15 DAH. Scale bar = 2 mm.

**Figure 3.**
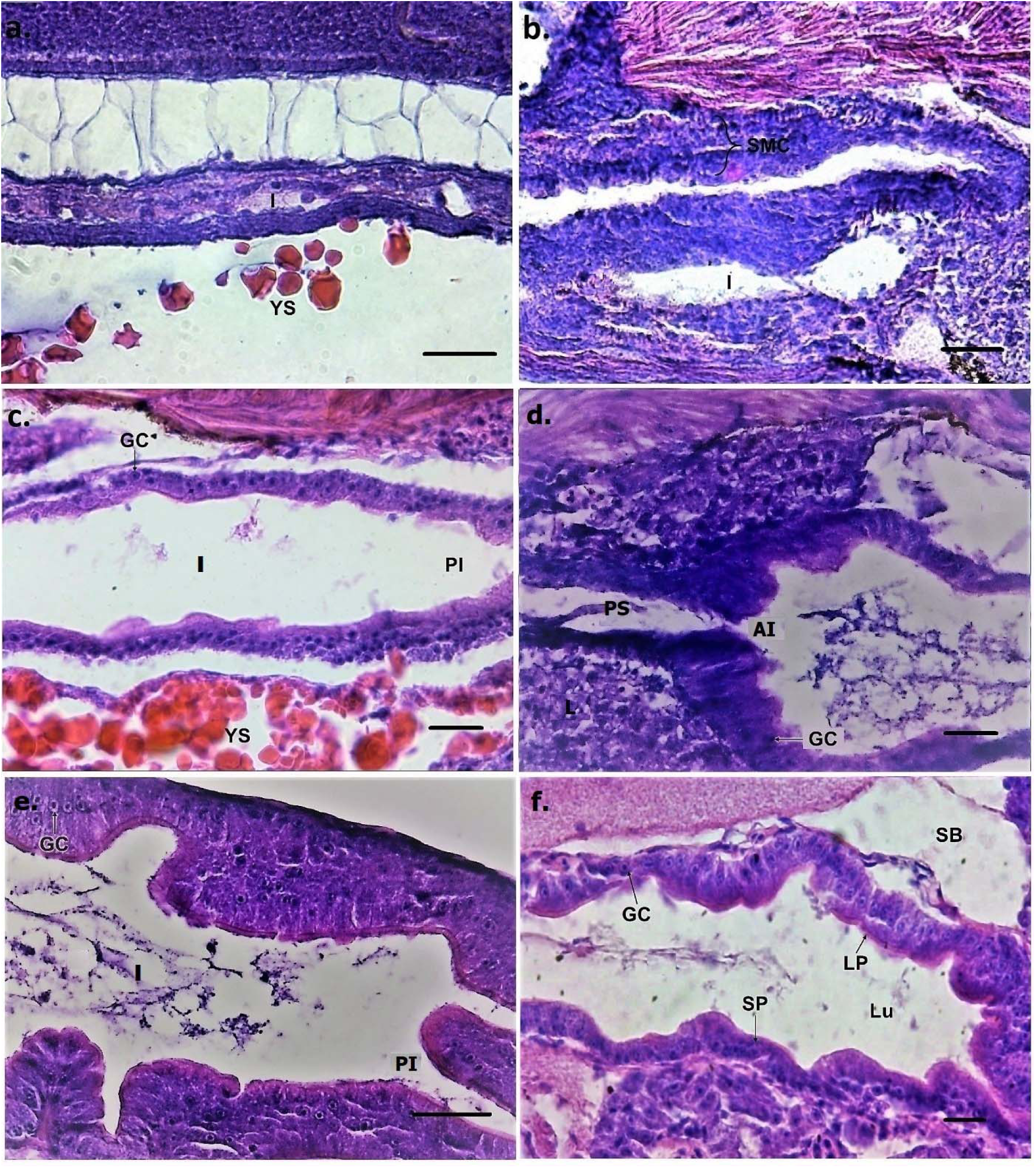
Histological on larval intestine. (a) Larval intestine at 1 DAH. Scale bar = 100 µm (b) Intestine started to elongate at 2 DAH. Scale bar = 10 µm (c) The posterior intestine and goblet cells were developed at 3 DAH. Scale bar = 100 µm (d) Posterior stomach was attached to the anterior intestine as early as 3 DAH. Scale bar = 100 µm (e) The goblet cells were dispersed at the intestine at 4-5 DAH. Scale bar = 10 µm (f) The digestive tract was fully developed with the layer of supranuclear protein and lipoproteins at 6 DAH. Scale bar = 10 µm. I, intestine; AI, anterior intestine; GC, goblet cells; IB, intestinal bulb; PI, posterior intestine; SB, swim bladder; SP, supranuclear protein; LP, lipoproteins; Lu, lumen; YS, yolk sac; SMC, simple muscle cells.

At 3 DAH, the digestive tract was connected to the mouth and anal openings that coincided with the start of exogenous feeding and followed by the coiling of the intestine. At this stage, the yolk sac was still not completely resorbed. The posterior intestine was elongated while the intestinal mucosa epithelium was composed of a single columnar layer and the goblet cells were surrounded by a thin layer of microvilli on their apical surface (Figure 3c). The stomach formation could be observed as early as 3 DAH. A few goblet cells began to differentiate and were observed amongst the epithelium in the same day. On the other hand, the mucosa was composed of undifferentiated simple cuboidal epithelium (Figure 3c). The number of goblet cells and rate of mucosal stratification increased with the age of the fish. The gill structure was formed from epithelium tissue with mucosa composed of simple cuboidal cells below the layer of the pharynx at 3 DAH (Figure 4a). In the same day, the intestinal tract began to differentiate from the stomach (Figure 3d) while the post-oesophagus swelled from the transition stratified epithelium to the columnar shape epithelial cells of the second portion of the oesophagus.

**Figure 4.**
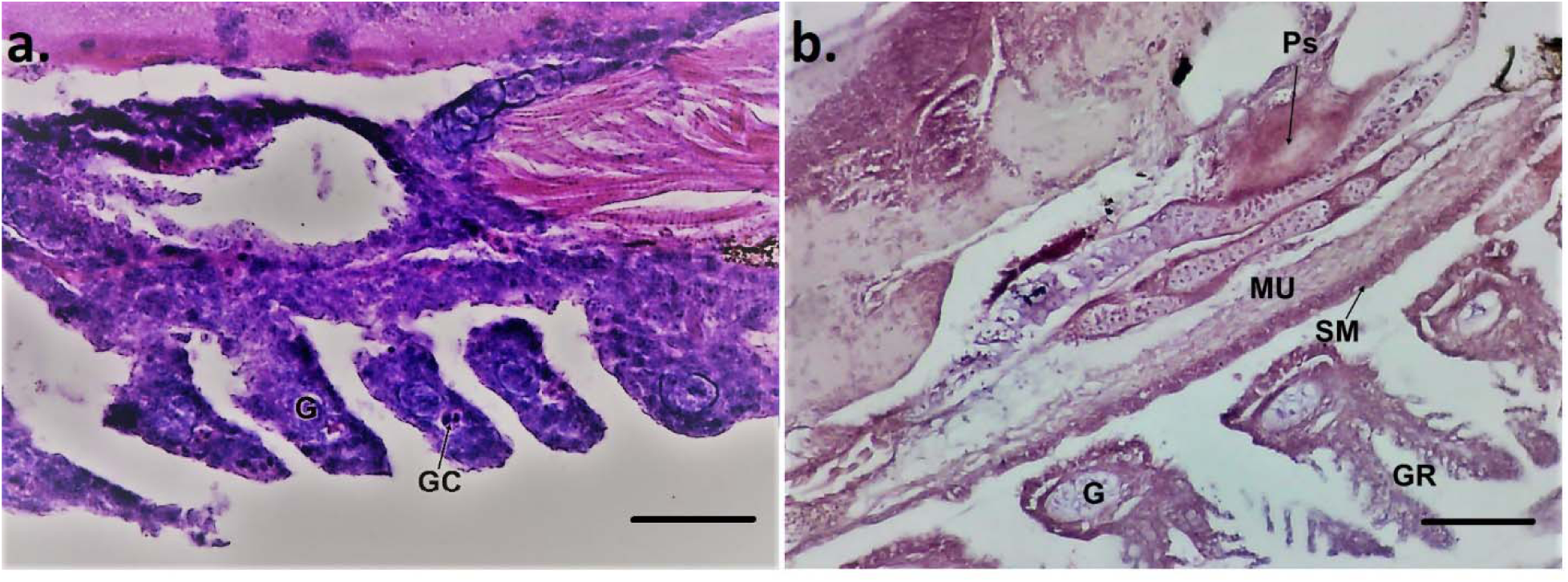
Histology of larval pharynx (a) Layer of pharynx at 3 DAH. Scale bar = 100 µm (b) pharynx attached to the muscularis and serous membrane at 6 DAH. Scale bar = 200 µm; G, gill; GC, goblet cells; GR, gill rakers Ps, Pseudobranch; MU, muscularis; SM, serous membrane.

The yolk sac was completely reabsorbed by the larvae at 4 DAH. By 5 DAH, the intestinal tract became more elongated from the anterior to the posterior intestine. Numerous goblet cells were visible throughout the oesophagus on the same day (Figure 3e). At this early stage, the posterior portion of the intestine had a higher number of mucous-secreting goblet cells than the anterior portion. Though it was established that the digestive tract comprised of six morphologically distinct regions namely buccal cavity, pharynx, oesophagus, stomach, anterior intestine and posterior intestine. The pharynx extended posteriorly past the gills where it was narrowed with the muscularis and serous membrane formed along oesophagus at 6 DAH (Figure 4b). The stomach began to increase in size and function at 6 DAH while the digestive tract became elongated towards the anus. First gastric glands could not be observed in this study. The layer of supranuclear protein and lipoprotein were well developed at the epithelium layer along the intestinal tract starting from 6 DAH (Figure 3f).

The length of digestive tract increased as fish grew, while the number of coils increased from 7 DAH with RGI value was 1.07 ± 0.06. The digestive tract was considered completely developed by 7 DAH in which the supranuclear protein was fully covered with lipoprotein throughout the outer layer tract. The goblet cells also became dispersed along the digestive tract especially in the buccal cavity and intestine by 7 DAH. Stomach bulge was formed in the middle of posterior stomach and anterior part of intestine at 10 DAH. Gastric gland was first observed at 13 DAH along with the presence of the pyloric caeca in the anterior part of stomach (Figure 5a). The number of folds and gastric glands continued to increase as the fish grew. The folds and gastric glands were observed in the stomach lined by a single cubical epithelium at 17 DAH (Figure 5b).

**Figure 5.**
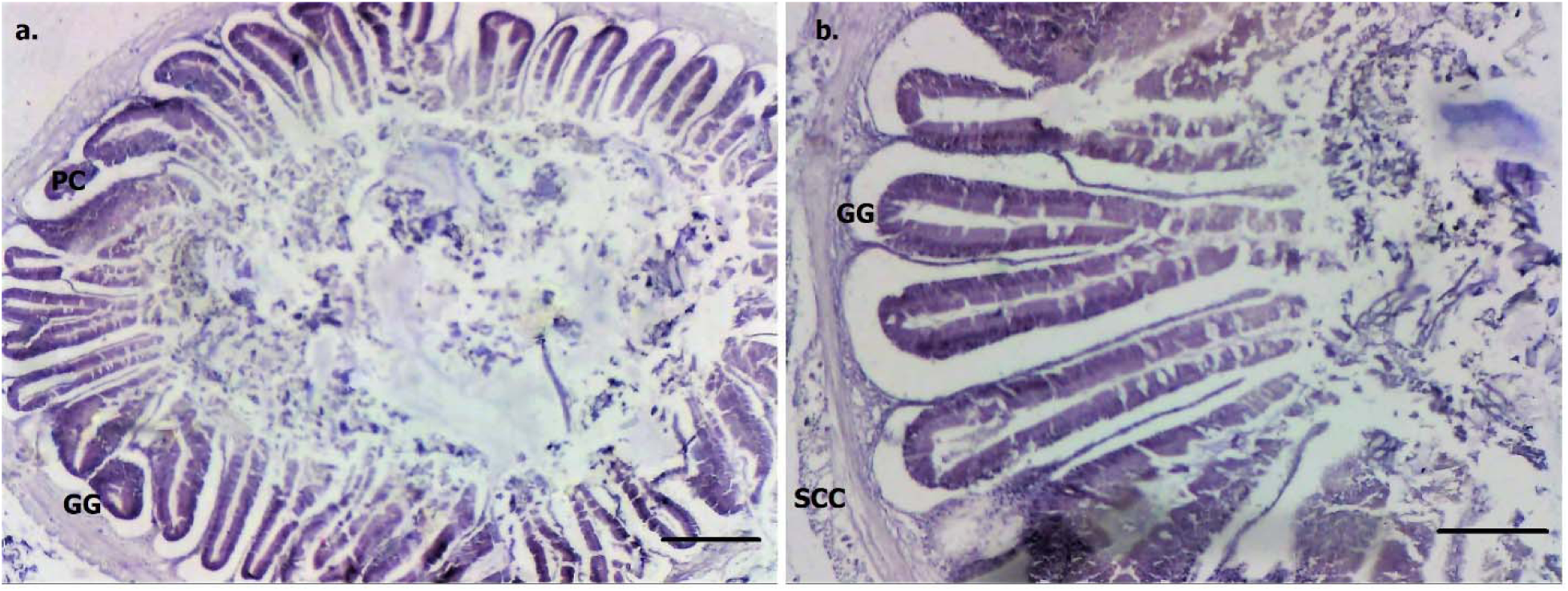
Histology of larval stomach (a) the presence of gastric glands at 13 DAH. Scale bar = 20 µm (b) Complete structure of folds and gastric glands at 17 DAH. Scale bar = 10 µm; GG, gastric gland, PC, pyloric caeca, SCC, simple cuboidal epithelium.

The structure of digestive tract looked similar to an adult fish by 23 DAH. No morphological change of digestive tract was observed beyond 23 DAH excepting for the gut length. The length of digestive tract continued to increase as the fish grew up to 3 times longer than the fish total length. Figure 6 shows the internal abdominal and digestive tract of hybrid Malaysian mahseer after a 2-month rearing.

**Figure 6.**
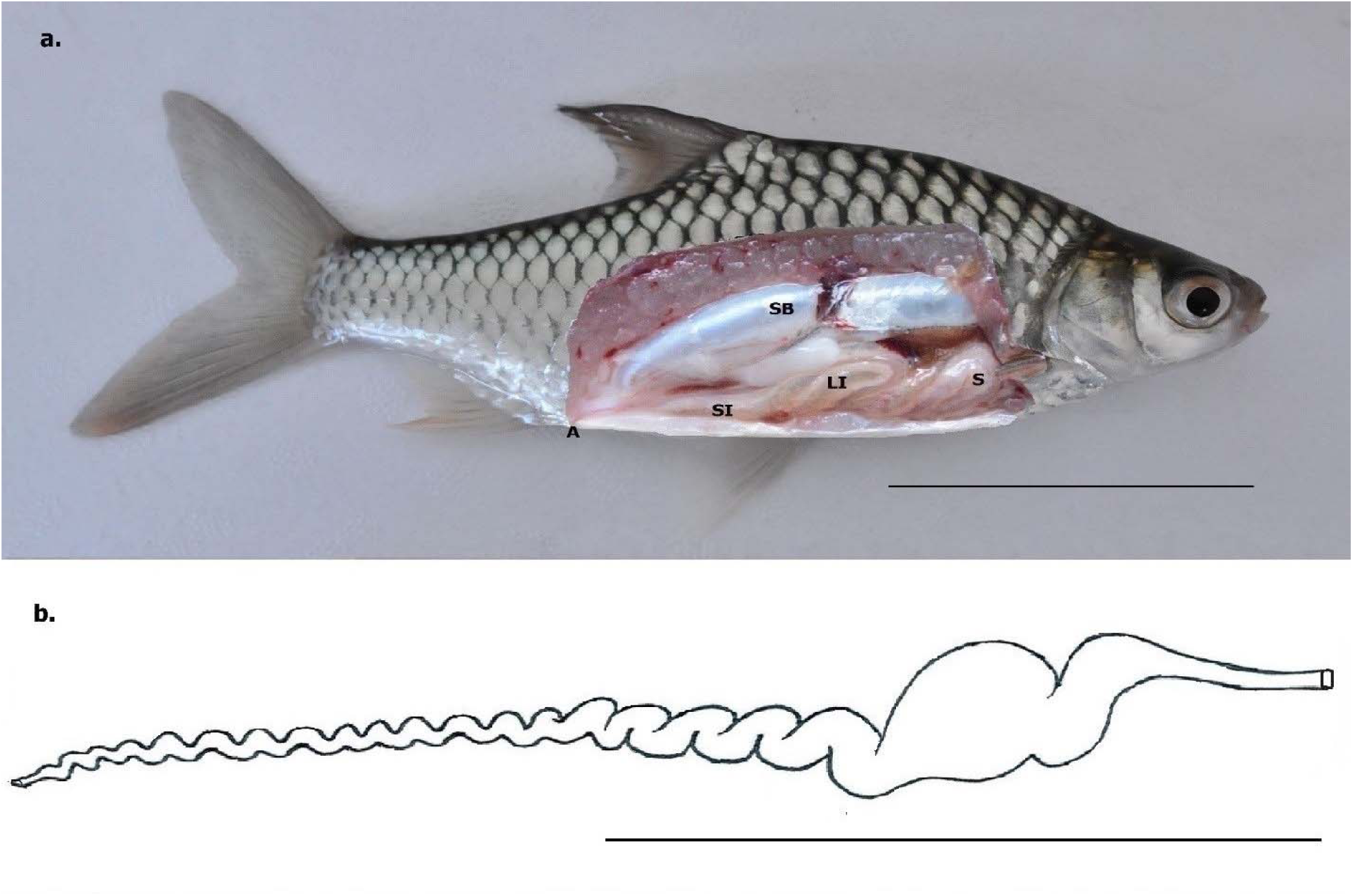
Digestive tract of 2-month old juvenile Malaysian mahseer hybrid a. Position of digestive tract in the abdominal cavity. b. Drawing of a dissected out digestive tract. Scale bar = 3 cm. A, anus; S, stomach; LI, large intestine; SB, swim bladder; SI, small intestine.

## 4. Discussion

Feeding ecology and behavior can be determined based on the gut length and relative length of the digestive tract (Das and Srivastava, 1979). It is evident that the value of RGI has a close relationship with the nature of fish diet (Dasgupta, 1988). The digestive tract length of herbivorous fishes is usually longer in comparison to carnivorous and omnivorous while the reverse is true for the number of digestive tract coils in carnivorous fish when compared to herbivorous and omnivorous fish (Das and Srivastava, 1979; Kamal, 1964). However, Dasgupta (2000) argued that digestive tract length may change according to the kind of diet provided. Generally, carnivorous and omnivorous fishes may require strong acid secretions and thick intestinal wall during their digestion process (Mohanta et al., 2009).

Fish with RGI above 0.8, 0.7-0.8 and under 0.7 considered to be herbivorous-omnivorous, carnivorous-omnivorous and carnivorous, respectively (Cahu and Zambonino Infante, 2001; Koundal et al., 2012). The RGI values in omnivorous fishes are smaller than those of herbivorous fishes since vegetables matters contain a high fiber content which require a longer digestion period. The RGI of hybrid Malaysian mahseer was considerably low in early larval stage. However the percentage increased with larval age. This hybrid fish had a similar pattern of digestive tract length and morphological functions with maternal fish, *B. gonionotus* (Basak et al., 2014; Mostakim et al., 2001). A juvenile hybrid mahseer had a RGI of about 2.5. Thus, the hybrid Malaysian mahseer could be considered as carnivorous fish during its early larval stage and changed to the herbivorous-omnivorous in a later grow-out stage. At a standard length of 16.4 mm, *B. gonionotus* has a RGI of 2.5 (Mostakim et al., 2001).

The presence of a short and straight digestive tract in newly hatched hybrid larvae was similar to findings of many earlier studies (Kamarudin et al., 2011; Ramezani-Fard et al., 2011; Teles et al., 2015). A *Iheringichthys labrosus* larva initially possesses a digestive tract with a flat anterior region, an elongated posterior and two intestinal folds (Kohn and Fernandes, 2011). The anterior region begins to differentiate into a stomach and the longer intestine with three intestinal folds in later stages (Makrakis et al., 2005). The onset of differentiation in the primordial cuboidal cells of the hybrid larval digestive tract at 2 DAH was similar to the larval Malaysian mahseer, *T. tambroides* (Ramezani-Fard et al., 2011). The proliferation of goblet cells in the posterior oesophagus has been found in yellowtail flounder at mid-larval stage (Baglole et al., 1997). The presence of goblet cells in the digestive tract of hybrid Malaysian mahseer larvae corresponded with the paternal fish, *T. tambroides*, where the first goblet cell appeared at 2 DAH (Ramezani-Fard et al., 2011). Mai et al. (2005) demonstrated that goblet cells are dispersed at the anterior oesophagus and the presence of lipid inclusion in the apical part of the epithelial cells of the large croaker teleost (*Pseudosciaena crocea*).

In this study, the number of goblet cells increased as the hybrid larva grew, whereby it started to disperse along anterior and posterior oesophagus by 5 DAH. Kozarić et al. (2008) stated that goblet cells help to secrete mucus for the purpose of protecting the alimentary canal against physicochemical damage and bacterial attack. This is important for the larval development as the mucus acts as a lubricant in food transportation along digestive tract as well as serving a saliva-like function. The presence of goblet cells during early larval development has been reported for *Silurus glanis* (Kozarić et al., 2008), *Solea senegalensis* (Ribeiro et al., 1999) and *Solea solea* (Boulhic and Gabaudan, 1992). The yolk is utilized for the embryonic and larval development as an endogenous feeding means with varying efficiency characteristics for different species (Parra and Yufera, 2001). The period of the complete yolk-sac absorption varies among species. Complete yolk sac absorption for *Hysibarbus malconi* and *Scophtalmus maximus* occurs by 2 DAH (Cunha and Planas, 1999; Ogata et al., 2010) while larvae of *Cyprinus carpio* (Haniffa et al., 2007), *T. tambroides* (Ramezani-Fard et al., 2011) and *Mystus nemurus* (El Hag et al., 2012) completely reabsorb their yolk sac by 3 DAH. The yolk sac was completely reabsorbed by hybrid Malaysian mahseer larvae at 4 DAH which was similar to captive red scorpionfish, *Scorpaena scorfa* (Maricchiolo et al., 2016) and silver perch, *Bidyanus bidyanus* (Sulaeman and Fotedar, 2017). Jafari et al. (2009) reported that a temperate carp, *Rutilus frisii kutum* takes a much longer (20 DAH) to completely absorb its yolk sac.

Exogenous feeding usually starts when the mouth is opened and the digestive tract is connected to both mouth and anus (Moteki et al., 2001; Teles et al., 2015). The present study showed that the larva of hybrid Malaysian mahseer began the exogenous feeding as early as 3 DAH which was similar to the same hybrid fish (Azfar Ismail et al., 2019), *Rutilus frisii kutum* larvae (Jafari, 2011), *B. gonionotus* (Basak et al., 2014) and *Puntius sarana* (Chakraborty et al., 2012). In contrast, the paternal *T. tambroides* larva starts exogenous feeding at 5 DAH with the appearance of clearly visible taste buds and goblet cells including secretion and stratified mucosal cells at the buccopharynx regions (Ramezani-Fard et al., 2011).

This present study revealed that the oesophaghus of larval hybrid mahseer was fully developed at 7 DAH which was similar to 5-7 DAH findings among several cyprinid fish (Ramezani-Fard et al., 2011). On the same day, the formation of a functional stomach of hybrid mahseer was completed. The intestinal bulb of this hybrid larvae was fully formed at 10 DAH. The results of the present study were comparable with the findings of Boulhic & Gabaudan (1992), Sarasquete et al. (1995), Ramezani-Fard (2011) and Kozarić et al. (2008). *T. tambroides* larvae are able to ingest, digest and absorb an artificial diet of 287 µm Ø from 7 DAH onwards with the completion of the morpho-histological development of digestive tract and the presence of supranuclear protein (Ramezani-Fard et al., 2011). These findings are aligned with the results of the present study where a functional stomach was observed at 6 DAH while the differentiation of digestive tract was completed at 7 DAH. This latter indicated that the larval hybrid Malaysian mahseer should be able to utilise an artificial diet as early as 7 DAH. In fact, the hybrid larvae were observed to readily consume and ingest the commercial larvifeed introduced in this study. El Hag et al. (2012) and Verreth et al. (1992) also reported that *Mystus nemurus* and *Clarias gariepinus* larvae have fully functional digestive tract at 5-7 DAH. Meanwhile, *Clarias lazera* larvae has a completely developed stomach by 4 DAH (Stroband and Kroon, 1981).

High protein and lipid contents are required in the diet of fish larvae because lipid helps to maximize protein sparing (Hasan, 2001). Larvae undergo a critical period when they are fed an artificial diet while having an inadequate digestive capability (Rønnestad et al., 2013). The processes in intracellular protein digestion may assist in the assimilation of macromolecular protein in the larvae because of the insufficient production of digestive enzymes and incomplete digestive capability (Gordon and Hecht, 2002; Sarasquete et al., 1995). Proximal intestine for most cyprinid fish usually absorbs fat rather than protein (Rombout et al., 1984). Supranuclear protein is the molecular protein found in the posterior intestine with the presence of intracellular protein digestion and pinocytotic absorption (Mai et al., 2005). Supranuclear protein appearance in the stomach within 8 DAH has been verified in some fish species (Baglole et al., 1997; Gordon and Hecht, 2002; Mai et al., 2005). Lipoproteins consist of non-polar lipids, primarily cholesterol esters and triglycerides in the central hydrophobic core that are attached to the lipid membrane which contains supranuclear protein (Feingold and Grunfeld, 2000). The lack of lipoprotein in the intestinal cells of larvae affects the accumulation of free lipid droplets in the cytoplasm. The improvement of lipoprotein synthesis and enterocyte development in hybrid mahseer larvae were similar to findings reported for *Sparus aurata* (Sarasquete et al., 1995) and *T. tambroides* (Ramezani-Fard et al., 2011).

## 5. Conclusion

Histo-morphological observations are essential in studying the development of digestive tract and feeding of larval fish. Exogenous feeding of hybrid mahseer larva began at 3 DAH which coincided with the opening of the mouth and anus, the connection of its alimentary tract to the anus and mouth, and the presence of developing stomach. The inclusion layer of supranuclear protein and lipoprotein in intestine, the appearance of a well-functioning stomach and the feeding observation strongly indicated that hybrid larva should be capable to accept and utilise an artificial diet by 7 DAH. Further studies should be conducted on the digestive enzyme development especially pepsin in the larval hybrid Malaysian mahseer to strengthen this notion.

## Acknowledgment

This study was funded by the Malaysian Government under Trans-disciplinary Research Grant Scheme (TRGS-KPM) through project no. 129831-138097 and Fundamental Research Grant Scheme (FRGS-KPM) through project no. 07-01-16-1795FR. The authors would like to thank the Institute of Bioscience of Universiti Putra Malaysia (UPM) for their cooperation and assistance.

